# Atypical Child-Parent Neural Synchrony Links to Children’s Psychopathological Symptoms

**DOI:** 10.1101/2022.03.30.486310

**Authors:** Haowen Su, Christina B. Young, Zhuo Rachel Han, Jianjie Xu, Bingsen Xiong, Jingyi Wang, Lei Hao, Zhi Yang, Gang Chen, Shaozheng Qin

## Abstract

Family emotional climate is fundamental to child’s emotional wellbeing and mental health. Negative family emotional climate may lead to heightened psychopathological symptoms via dysfunctional child-parent interactions. Single-brain paradigms have uncovered changes in brain systems and networks related to negative family environments, but how neurobiological reciprocity between child and parent brains is associated with children’s psychopathological symptoms remains unknown. In study1, we investigated the relationship between family emotional climate and children’s psychopathological symptoms in 395 child-parent dyads. In study2, using a naturalistic movie-watching functional magnetic imaging technique in a subsample of 100 children and parents, we investigate the neurobiological underpinnings of how family emotional climate is associated with children’s psychopathological symptoms through child-parent neural synchrony. Children from negative family emotional climate experienced more severe psychopathological symptoms. We revealed significantly higher inter-subject correlations in the dorsal and ventral portions of the medial prefrontal cortex (mPFC), and greater concordance of activity with widespread brain regions critical for socioemotional skills in child-parent than child-stranger dyads. Critically, negative family emotional climate was associated with decreased inter-subject functional correlation between the ventral mPFC and the hippocampus in child-parent dyads, which further accounted for higher children’s internalizing symptoms especially for anxious and depressed aspects. Family emotional climate might transmit into the brain of parent-child dyads, which may associate with child development outcomes. The present study identified that child-parent vmPFC-hippocampal circuitry is linked to children’s psychopathological symptoms. Our findings suggest a neurobiological mechanism of how negative family emotional climate affects children’s psychopathological symptoms through altered child-parent neural synchrony.

## Introduction

Family has been considered the primary socialization contexts for children, in which children begin to learn how to appropriately express and regulate their emotions through observing and modeling their parents’ emotional behaviors (Eisenberg, 2020; McCoy & Raver, 2011). Family emotional climate, including family negative and positive emotion expression and specific emotion-related parenting, have great importance of shaping children’s emotional wellbeing and mental health(Speidel et al., 2020). Research has indicated that family emotional climate, especially its negative aspect, can cause and maintain children’s various psychopathological symptoms(Gong et al., 2021; Rea et al., 2020; Teicher et al., 2016).

Psychosocial views suggest that negative family emotional climate may lead to children psychopathology symptoms through derailing of the coordination of moment-to-moment behaviors between child-parent dyads (Feldman, 2020b; Morris et al., 2018). Indeed, previous studies have demonstrated that negative family environments compromise the effectiveness of reciprocal interactions in child-parent dyads (Hoyniak et al., 2021a; Tarullo et al., 2017). Children who experienced maladaptive family interaction with parents are prone to develop psychopathological symptoms later (Feldman, 2012) in life (Feldman, 2007; Quiñones-Camacho et al., 2021). According to bio-behavioral synchrony model, the coordination of children-parent interaction contains various components such as behavior, autonomic, hormones, and brain, which are closely associated with child developmental outcomes(Feldman, 2012). Although well documented in behavioral (e.g.,Thomassin & Suveg, 2014)and physiological studies (e.g., Davis et al., 2018), the underlying neurobiological mechanisms of how family emotional climates associate with child psychopathological symptoms via altered reciprocal responses across child-parent brains remain largely unknown.

With the concentration on cross-brain associations, the Extended Parent–Child Emotion Regulation Dynamics Model (Ratliff et al., 2022) proposed that parent-child brain-to-brain concordance is a key mechanism linking family system (e.g., family emotional climate) with child psychopathological symptoms. Guided by this framework, researchers have conducted studies to understand how indicators of family environment associated with parent-child brain-to-brain concordance (e.g., interbrain synchrony, shared neural similarity), and how the interbrain concordance is associated with children’s psychopathological symptoms. Using functional near-infrared spectroscopy hyperscanning techniques, recent studies have demonstrated neural synchrony in the prefrontal cortex (PFC) across brains critical for child-parent reciprocal interactions including cooperation (Reindl et al., 2018a), joint attention and smiling(Piazza et al., 2020). Family risk factors including parenting stress (Azhari et al., 2019), maternal stress (Nguyen et al., 2020), anxious attachment (Azhari et al., 2020), and sociodemographic risks (Hoyniak et al., 2021b) could diminish child-parent shared neural response in the PFC. Moreover, disrupted child-parent prefrontal synchrony was associated with children’s poor emotion regulation(Reindl et al., 2018a), heightened irritability (Quiñones-Camacho et al., 2020), and salient autism spectrum disorder (ASD) symptoms among children with ASD(Wang et al., 2019). These findings provide initial evidence to suggest that family risk factors may alter child-parent brain-to-brain concordance, thus impeding children’s socioemotional ability in development.

In addition to examining brain-to-brain synchronization during parent-child interaction, the transmission of such shared representations may exhibit shared neural activity patterns between child and parent brains in response to any related events even without real-time interactions. Children learn social and emotional skills through dyadic interactions with their parents, which can help form and maintain shared neural representations of socioemotional experiences and knowledge in long-term memory and then contribute to their development (Feldman, 2007a; Fiske & Taylor, 2013). These processes require multiple brain regions and systems to interact and exchange information (Babiloni & Astolfi, 2014). Empirically, researchers have applied a sequential dual-brain fMRI paradigm (scans a single participant each time engaged in a shared task with two identical sessions to investigate the shared/different brain responses) to examine the psychological function of shared neural responses(Lee et al., 2017, 2018). Lee et al. (2018) indicated that mother–child dyads with lower levels of family connectedness showed less similar neural response patterns in anterior insular and dorsal anterior cingulate cortex (dACC), which then accounted for adolescent heightened perceived stress level during stress tasks. In addition, lower levels of resting state whole-brain connectome similarity were associated with children’s poorer emotional competence, which might increase the risk of suffering psychopathological symptoms (Lee et al., 2017). However, existent studies about shared neural responses did not identify the role of PFC, especially the medial prefrontal cortex (mPFC), which cannot be examined via functional near-infrared spectroscopy and electroencephalogram due to the lack of space resolution and detecting deep brain regions.

The medial prefrontal cortex (mPFC), a core node of the social and emotional networks, is recognized to play a critical role in the transmission of shared socioemotional schema involving knowledge, values and beliefs across individuals (Krueger et al., 2009a; Roy et al., 2012a).

Indeed, recent studies demonstrate that the mPFC works in concert with the hippocampus allowing individuals to learn and accumulate necessary socioemotional knowledge and experience through social interactions(Hiser & Koenigs, 2018; Yeshurun et al., 2021).

Methodologically, mapping brain-to-brain concordance has the potential to advance our understanding on how shared socioemotional representations across brains arise from dyadic interactions between children and parents (e.g., Reindl et al., 2018). In particular, recent studies have utilized naturalistic movie watching fMRI to detect how individuals process and understand the complex socioemotional world based on our internal mental schema including long-term memory, emotion and prior belief. This paradigm lends itself to enabling inter-subject correlation (ISC) and inter-subject functional connectivity (ISFC) analyses to identify brain activity concordance in cortical and subcortical regions among subjects, and their related functional circuits during movie watching (Hasson et al., 2004a; Simony et al., 2016). Therefore, these approaches compute functional activity concordance across brains, and are emerging as a powerful tool for exploring brain function and their concordance across individuals that allows us to identify: 1) shared neural responses across child-parent brains during movie watching, 2) reactivity of related socioemotional knowledge in long-term memory in response to boundary and non-boudary events, 3) child-parent dyadic brain predictors underlying the effects of family emotional climate on children’s psychopathological symptoms. Based on the Extended Parent– Child Emotion Regulation Dynamics Model (Ratliff et al., 2022), we hypothesized that child-parent dyads would exhibit increased brain-to-brain concordance in the mPFC and related functional circuits compared to control dyads (i.e., child-stranger dyads), and that amount of concordance would mediate the association between negative family emotional climate and children’s psychopathological symptoms.

To test the above hypotheses, we conducted two separate studies integrating behavioral assessments of family emotional climate and children’s psychopathological symptoms, as well as dyad-based analysis of movie-watching fMRI data in child-parent and child-stranger dyads. In study 1, we investigated how family emotional climate related to children’s psychopathological symptoms including internalizing and externalizing problems in 395 child-parent dyads (**Figure. 1a**). We assessed family emotional climate using a well-validated scale that characterizes how often positive and negative emotions are expressed in a family (Halberstadt et al., 1995a). Internalizing and externalizing symptoms were measured by the widely used child behavior checklist (CBCL) (Achenbach, 1991). In study 2, we used a child-friendly naturalistic movie-watching paradigm in an fMRI experiment known for its ecological validity (Vanderwal et al., 2019) to measure child-parent shared brain responses in 50 child-parent dyads (**Fig. 1b**). Brain-to-brain concordance metrics in response to movie watching were assessed through ISC and ISFC methods. Given the correlational nature of dyad-based brain concordance, we used an optimized linear mixed effect model with crossed random effects to account for the complex data hierarchical structure(Chen et al., 2017) (**Figure. 1c**). Mediation analyses were used to examine if child-parent shared brain response accounts for the direct and/or indirect effects of negative family emotional climate on children’s internalizing and externalizing symptoms.

**Figure 1.**
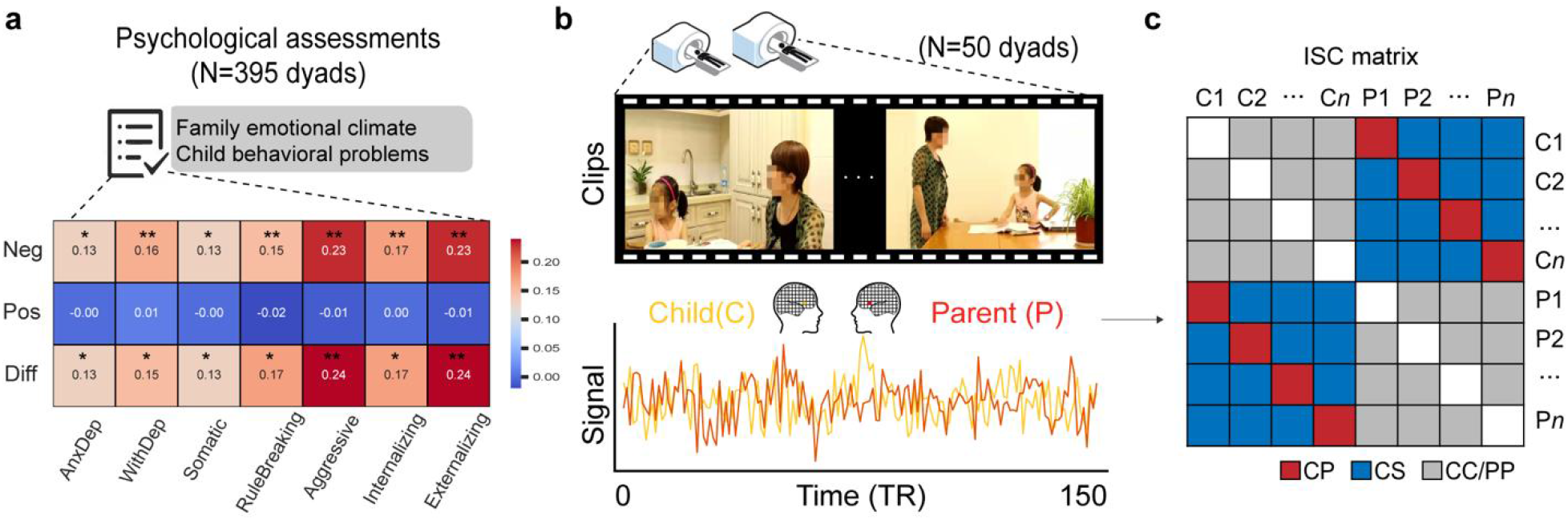
An Illustration of Experimental Design and Inter-Subject Correlation Analysis (ISC) *Note*. a. Correlations of negative (‘Neg’) and positive (‘Pos’) family emotional climate with children’s internalizing and externalizing problems respectively. The difference between the negative (top row) and positive (middle row) family emotional climate is shown with strong statistical evidence. b. Representative frames of a 6-min movie of a 7-year-old girl and her mother used as the fMRI paradigm. An illustration of voxel-wise ISC between two time series of a child-parent dyad for each voxel of the grey matter mask. c. A matrix represents pairwise correlations among child and parent subjects, resulting in child-parent dyads (CP, red), child-stranger controls (CS, blue), child-child & parent-parent pairs (CC/PP, grey). Each cell represents a ISC value. *, *q* < 0.05; **, *q* < 0.01.

## Methods and Materials

### Participants

A total of 446 families participated in Study 1. After removing participants with missing values (missing > 20%) in behavioral measures, data of 395 child-parent dyads (children: mean ± S.D. = 9.35 ± 1.64 years old, range 6.28-12.51, 50% boys; parents: mean ± S.D. = 37.42 ± 4.62 years old, range = 27-57, 33% father) were analyzed in the correlation between family emotional climate and child’s psychopathological symptoms. All participants had normal or corrected-to-normal vision, and no one reported a history of psychiatric or neurobiological disorders. A subset of 50 child-parent dyads completed fMRI scanning while watching a negative movie clip in Study 2. Nine child-parent dyads were excluded due to children’s head motion with mean framewise displacement larger than 0.5 mm during movie watching scanning. The final sample consists of 41 child-parent dyads (children: mean ± S.D. = 10.15 ± 1.41 years old, range 7-12, 46.3% boys; parents: mean ± S.D. = 38.91 ± 5.34 years old, range=29-49, 24.4% father). Six child-parent dyads were further excluded for the mediation analyses due to the incomplete family emotional expressiveness questionnaires. In the rest sample, after excluding subjects with mean framewise displacement larger than 0.5 mm, data of 25 pairs of children and parents were analyzed in the ISC and ISFC (children: mean ± S.D. = 7.8 ± 1.33 years old, range 7.80-12.26, 40% boys; parents: mean ± S.D. = 39.31 ± 5.51years old, range=29 - 49, 16% father). All subjects provided written informed consent before their participation and received monetary compensation. The study was approved by the Institutional Review Board (IRB) of the local institute.

### Materials and Movie Watching

A 6-minute video unfamiliar to all participants was shot for this study, which showed a 7-year-old girl arguing with her mother (**Figure. 1b**). Critical moments are provided in *Table S1*. Children and their parents underwent fMRI when separately viewing the video. This video was played with no sound/subtitles to mitigate potential confounds in auditory perception and language comprehension.

### Psychological assessment

#### Psychopathological Symptoms

Children’s psychopathological symptoms were assessed by the parent-reported CBCL scores based on parent surveys (Achenbach, 1991), including anxious/depressed syndrome withdrawn/depressed syndrome, somatic, score and rule-breaking, internalizing and externalizing score. The Chinese version of the CBCL has been used widely(Crijnen et al., 1999). The internal consistency of the parent-reported CBCL scales in the present study was α = 0.81 for internalizing problems and α = 0.84 for externalizing problems. The raw scores of all sub-scales were used in all analyses.

#### Family Emotional Climate Assessment

Family emotion climate was measured by the Family Expressiveness Questionnaire (EFQ) (Halberstadt et al., 1995b). Parents reported how often positive and negative emotions were expressed in their family on a 9-point Likert scale. Coefficient alphas in the present sample were 0.89 and 0.85 for the positive and negative subscales.

#### Family Socioeconomic Status Assessment

Family socioeconomic status was measured by a self-report family background questionnaire that used 10- and 6-point scales to assess the education and monthly income of each parent respectively. To form a composite SES score for each parent, the income and education scores were first divided into individual z scores for each parent, which were then averaged.

### Brain Imaging Data Acquisition

Whole-brain images were acquired from Siemens 3.0 T scanner (Siemens Magnetom Trio TIM, Erlangen, Germany), using a 12-channel head coil with a T2*-sensitive echo-planar imaging (EPI) sequence based on blood oxygenation level-dependent (BOLD) contrast. Thirty-three axial slices (4 mm thickness, 0.6 mm skip) parallel to the anterior and posterior commissure (AC-PC) line and covering the whole brain. Each participant’s high-resolution anatomical images were acquired through three-dimensional sagittal T1-weighted magnetization-prepared rapid gradient echo (MPRGE) with a total of 192 slices. Detail parameters and procedures are provided in supplementary information (SI).

### Brain Imaging Data Analysis

#### Preprocessing

Based on previous studies on ISC pre-processing pipelines (Nastase et al., 2019), brain images were preprocessed using statistical parametric mapping (SPM12). Images were corrected for slice acquisition timing and realigned for head motion correction. Subsequently, functional images were co-registered to each participant’s gray matter image segmented from corresponding T1-weighted image, then spatially normalized into a common stereotactic MNI space and resampled into 2-mm isotropic voxels. Images were smoothed by an isotropic 3D gaussian kernel with 6-mm full-width half-maximum (FWHM). The preprocessed images were regressed on a set of nuisance covariates (i.e., motion parameters, the average signal of white matter and cerebrospinal fluid) and 140-sec high-pass filtered using toolbox Nilearn version 0.6.2. Finally, the first 5 volumes and the last 5 ones were removed to minimize stimulus onset and offset effects and the data were z-scored overtime.

#### Inter-Subject Correlation (ISC) and Statistical Analysis

Whole-brain ISC maps during movie watching were computed for all possible pairs of 41 participants in a gray-matter mask using BrainIAK’s ISC function(Kumar et al., 2020) (Figure. 1b). The ISC maps were submitted to further statistical analyses to identify brain regions that show synchronous (shared) neural response across the whole sample. We adopted a linear mixed-effects (LME) model using a crossed random-effects formulation which can accurately interpret the ISC data’s correlation structure (Chen et al., 2017). Participants’ gender, age, and scanning sites were treated as covariates of no interest in the LME model. False discovery rate (FDR) correction was used to correct multiple comparisons (Benjamini & Hochberg, 1995). Next, we used a two-group formulation of the LME model with three covariates (age, gender, and site) to identify whether there were voxels that were more synchronous in child-parent dyads than child-stranger pairs. Child-parent dyads were defined by pairing a child and their own parent, and child-stranger dyads were generated by pairing a child with all parents except their own parent. 3dClustSim module of AFNI was used to correct multiple comparisons, with a voxel-wise *p* < 0.001, cluster-wise *α* < 0.05.

#### Inter-Subject Functional Connectivity (ISFC) Analysis

ISFC analysis was implemented to identify movie-evoked functional connectivity across participants. We used a seed-based ISFC approach by computing the correlation of a given seed’s [i.e., 6-mm sphere of the peak voxel at MNI coordinate (2, 38, −18) in the vmPFC and (0,56,12) in the dmPFC] time series in one participant with every other voxel’s time series in another participant. The computation of ISFC produced two asymmetric matrices for *r*_(VMPFCsubject1,Ysubject2)_ and *r*_(VMPFCsubject2,Ysubject1)_. We then computed the average correlation, which was treated as the ISFC value between each participant pair where *r* represents Pearson’s correlation and Y represents the time series of each given voxel from participants. Likewise, the LME was used to determine which brain regions showed higher coordination with vmPFC and dmPFC in child-parent pairs than child-stranger pairs. False discovery rate (FDR) was used to correct multiple comparisons ^42^.

#### Intra-Subjective Functional Connectivity (FC) Analysis

To verify that child-parent vmPFC-hippocampal ISFC played a unique role in the relationship between negative family emotional climate and children’s internalizing problem, we also examined whether the single brain’s vmPFC-seed FC is associated with negative family emotional climate and children’s internalizing problem. We examined single brain FC in children’s and parent’s brains, and ran multiple regressions with negative family emotional climate and children’s internalizing problems, as separate regressors, predicting vmPFC-seed function connectivity. Other settings are identical for above ISFC.

#### Meta-Analytic Decoding with Neurosynth

The neurosynth allows us infer the psychological domains involved in brain map of the shared vmPFC-circuits in child-parent dyads. Specifically, we correlated the child-parent thresholded vmPFC and dmPFC ISFC map (FDR *q* < 0.05) to the topics map of 15 general psychological domains involving a range of possible brain processes during movie viewing using the Neurosynth’s python notebook ([https://github.com/neurosynth/neurosynth]; commit version 948ce7).

#### Mediation Analysis

Mediation analysis was performed using the Mediation Toolbox developed by Tor Wager’s group (https://github.com/canlab/MediationToolbox). Prior to the mediation analysis, average values representing intersubject functional connectivity strength with vmPFC and dmPFC were extracted from 13 significant clusters and 2 significant clusters identified in the above linear mixed model to examine the correlation with negative family emotional climate using FDR corrections (67) to control the false-positives. Next, a mediation model was constructed to investigate the mediating pathways between negative family emotional climate, shared vmPFC-hippocampus ISFC strength, and children’s anxious/depressed and internalizing problems. The indirect or mediated effect was tested by a bias-corrected bootstrapping method (n = 10000 resamples). All statistical tests here are two-tailed and pass the FDR correction. More details are provided in SI.

#### Event Boundary Analysis

The correlation between hippocampal, vmPFC responses and event rating. Event boundaries were collected by an independent group of 20 adult raters (10 males) who watched this 6-minute silent video. The rates were asked to press a key at the end of one meaningful event and the beginning of another.In line with Reagh’s studies (68), we include the boundary timepoints and non-boundary timepoints of this video. Boundaries timepoints are agreed by at least half of the samples, and we find a total of 10 event boundaries of the time series. We also add the same number of non-boundary time points compared to the event boundaries. The boundary and non-boundary time points were next convolved with a canonical HRF to obtain the boundary and non-boundary timeseries. Then we correlated each participant’s hippocampal and vmPFC timeseries with the event boundary and non-boundary timeseries derived from the independent raters. Finally, we examined whether children’s hippocampal and vmPFC response to boundary and non-boundary timeseries were correlated with their parents’. All statistical tests here are single-tailed and pass the FDR correction.

## Results

### Negative Family Emotional Climate Linked to Children’S Internalizing/Externalizing Symptoms

First, we examined how family emotional climate, including positive and negative components was associated with children’s internalizing and externalizing symptoms in Study 1. Pearson’s correlation analyses revealed that negative family emotional climate was associated with more severe children’s internalizing symptoms (*r*_395_ = 0.17, *q* < 0.001, 95% CI = [0.08, 0.26]), including anxious/depressed (*r*_395_ = 0.13, *q* = 0.024, 95% CI = [0.03, 0.22]), withdrawn/depressed (r395 = 0.16, *q* = 0.005, 95% CI = [0.07, 0.25]), and somatic problems (*r*_395_ = 0.13, *q* < 0.001, 95% CI = [0.04, 0.22]), as well as externalizing symptoms (*r*_395_ = 0.23, *q* < 0.001, 95% CI = [0.14, 0.32]), including aggressive (*r*_395_ = 0.24, *q* < 0.001, 95% CI = [0.07, 0.24]) and rule-breaking behaviors (*r*_395_ = 0.15, *q* = 0 .001, 95% CI = [0.14, 0.32]) (*all q values were FDR corrected*) (**Figure. 1a**). There were no reliable associations of positive family emotional climate and children’s internalizing and externalizing symptoms (all *rs*395 < 0.01, *qs* > 0.70). Further tests for Fisher’s z-transformed correlation coefficients revealed statistically stronger correlations with negative than positive family emotional climate (all *Zs* > 1.95, *qs* < 0.05, FDR corrected). Notably, the positive associations of negative family emotional climate with children’s internalizing and externalizing symptoms remained statistically significant after controlling for child’s and parent’s age, gender, and socioeconomic status (*Supplementary Figure. S1*). These results indicate that children from negative family environment exhibit more severe internalizing and externalizing symptoms.

### Increased Inter-Subject Correlation in Vmpfc and Dmpfc during Movie Watching for Child-Parent Dyads

Next, we identified shared patterns of temporal neural activity in response to viewing a negative movie across brains. The ISC maps were computed to represent shared brain activity by correlating time series of the same voxel across participants (**Figure. 2a**). A linear mixed-effects (LME) model was conducted for ISC maps collapsing across children and parents to identify brain regions showing inter-subject synchrony during movie watching. This analysis revealed significant clusters in unimodal and transmodal association areas (**Figure. 2b**, *q* < 0.05 FDR-corrected). This pattern of results is consistent with ISC data from previous fMRI studies(Finn et al., 2018; Hasson et al., 2004b).

**Figure 2.**
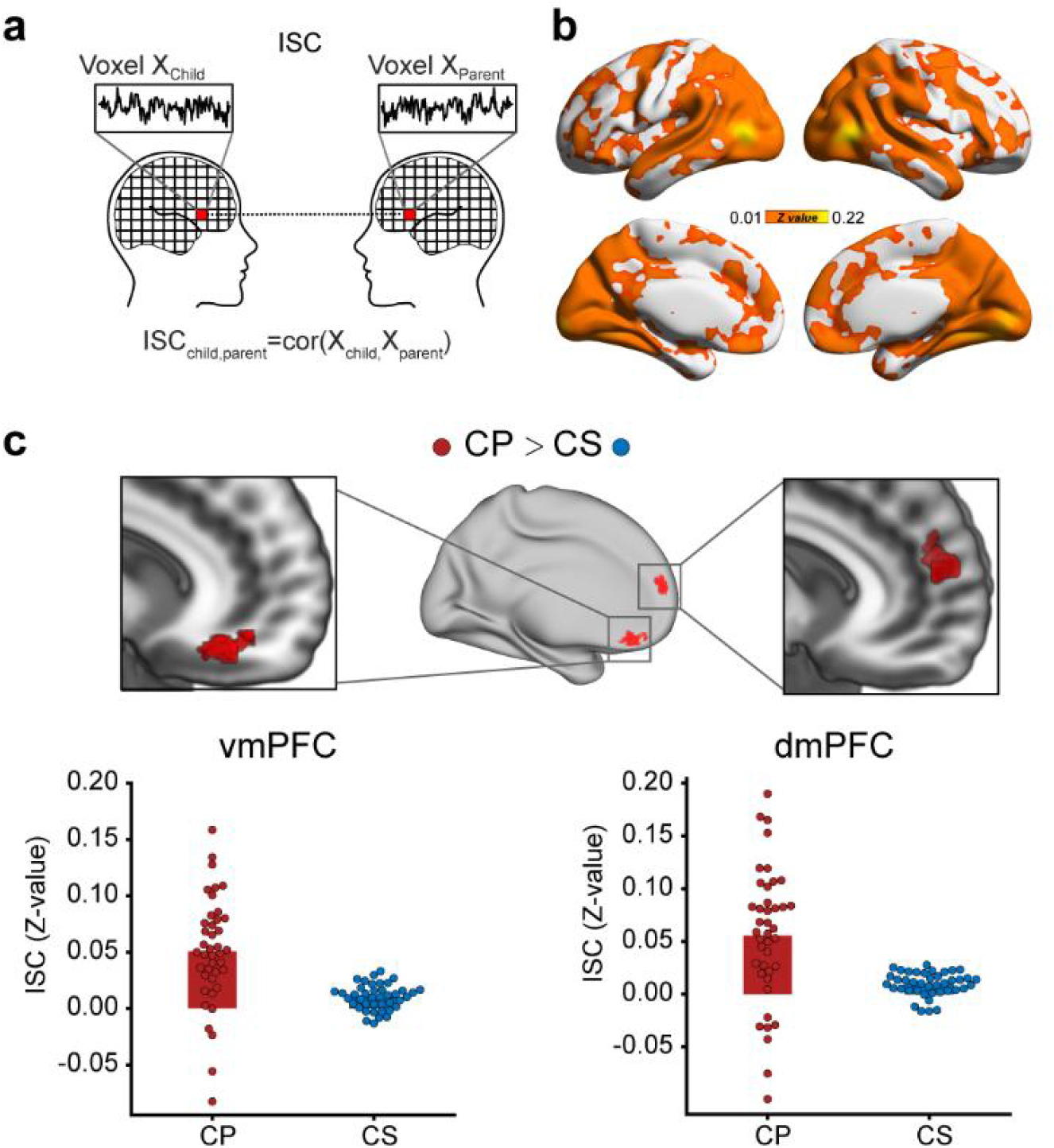
Primary Results from Inter-Subject Correlation (ISC) Analysis. *Note*. a. An illustration of inter-subject correlation (ISC) between time series of a given voxel in each child and his/her parent’s brain. b. Brain regions show statistically significant ISC during movie watching in general, with most prominent effect in the visual cortex followed by frontal, temporal, and parietal cortices. Statistically significant clusters were thresholded using *q* < 0.05 FDR corrected. The color bar represents Fisher’s Z-value. c. Representative views of the vmPFC and dmPFC showing stronger inter-subject synchronized activity (ISC) in child-parent dyads as compared to child-stranger controls. Statistically significant clusters were derived from a contrast between child-parent (CP) dyads and child-stranger (CS) controls, with a voxel-wise threshold *p* < 0.001 (two-tailed) combined with cluster-level threshold significance level α of 0.05.

We conducted dyad-based analysis using ISC maps between each child and their parent in comparison to each child and a stranger’s parent as a control. We implemented an optimized LME model with crossed random effects (Chen et al., 2017), and examined brain systems showing shared temporal neural responses during movie watching that unique to child-parent dyads relative to child-stranger controls. This analysis (**Figure. 2c**, *Supplementary Table S3*) revealed significant clusters (voxel-wise *p* < 0.001 two-tailed, cluster-wise significance level *α* < 0.05) in the vmPFC [peak MNI coordinate at (2,38, −18); cluster size *k* = 116 voxels] and the dorsal mPFC (dmPFC) [peak at (0, 52, 12), *k* = 122 voxels]. There were no surviving significant clusters when examining greater activity in child-stranger versus child-parent dyads. To verify whether this effect is specific to movie stimulus, we also performed parallel analysis for resting state fMRI data, and there were no statistically reliable ISC effects in the vmPFC and dmPFC.

### Increased Child-Parent Vmpfc Connectivity with Social and Emotional Systems during Movie Watching

Given that mPFC-centric circuitry is implicated in human emotion and social cognition (Krueger et al., 2009b; Lieberman et al., 2019), we used the vmPFC and dmPFC clusters identified above as separate seeds to perform inter-subject functional connectivity analyses. The LME model for vmPFC-seeded ISFC map was examined to identify functional circuits showing higher inter-subject connectivity in child-parent versus child-stranger dyads (Figure. 3a). This analysis revealed statistically significant clusters in widespread regions in the frontal, temporal and occipital lobes, including the hippocampus [peak MNI coordinates (−16, −30, −8)], amygdala [peak MNI coordinates (18,2, −16)], and fusiform gyrus [peak MNI coordinates (−32, −48, −8)], FDR *q* < 0.05 (**Figure. 3b-d**, Supplementary *Table S4*). Parallel analysis for dmPFC-seeded ISFC maps revealed that child-parent dyads exhibited higher connectivity with the angular gyrus [peak MNI coordinates (−46, −64, 24)] and medial prefrontal gyrus [peak MNI coordinates (0,54,14)] (*Supplementary Figure. S2a, Table S4*) than child-stranger dyads, FDR *q* < 0.05. To verify whether this effect is specific to movie watching, we also performed parallel analysis for resting-state fMRI data, and there were no any reliable ISFC effects in child-parent dyads compared to child-stranger controls.

**Figure 3.**
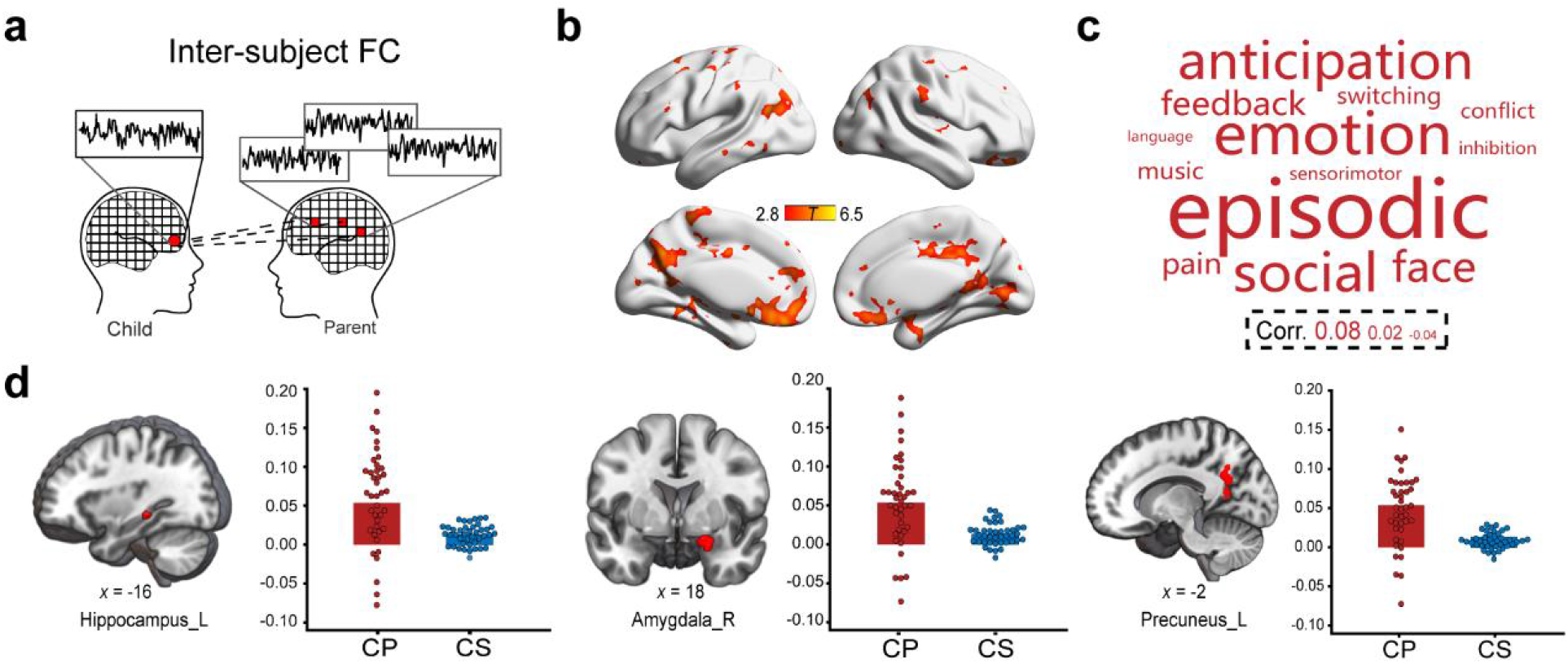
Results from Inter-Subject Functional Connectivity (ISFC) Analysis. *Note*. a. An illustration of seed-based ISFC that involves computing the correlation between a seed’s time series in a child brain and all other voxel’s time series of his/her parent brain. b. Compared to child-stranger controls, child-parent dyads showed stronger ISFC of the vmPFC with inferior frontal gyrus, middle cingulum gyrus, precuneus, fusiform, hippocampus and middle occipital gyrus (df = 77, *q* < 0.05 FDR corrected). c. Word cloud depicting commonly used terminology associated with regions showing vmPFC connectivity. d. Representative slices of significant clusters in the hippocampus, amygdala and precuneus that show stronger ISFC in child-parent dyads than child-stranger control dyads.

We then used a meta-analytic decoding approach based on a widely used Neurosynth platform (Yarkoni et al., 2011) to determine psychological functions of the above clusters that showed higher inter-subject connectivity with the vmPFC and dmPFC in child-parent than child-stranger dyads. This analysis revealed that child-parent shared vmPFC-based connectivity patterns with widespread regions that are implicated in episodic memory, emotion, and social functions **(Figure. 3c)**, whereas dmPFC-based connectivity did not exhibit a connectivity pattern implicated in these functions (*Supplementary Figure. S2c*). These results indicate higher vmPFC connectivity with social and emotional systems in child-parent dyads than child-stranger controls.

### Reduced Child-Parent Vmpfc Connectivity with the Hippocampus Mediates the Association between Negative Family Emotional Climate and Children’S Internalizing Symptoms

Given our central hypothesis that neural mechanisms link negative family emotional climate and psychopathological symptoms, we further investigated how negative family emotional climate shapes inter-subject correlation of brain activity and connectivity during movie watching in child-parent dyads, which was then associates with children’s psychopathological symptoms. Brain-behavior association analyses were conducted for ISC and ISFC metrics of the vmPFC and dmPFC. With these metrics, we found that family emotional climate was significantly correlated with lower child-parent shared vmPFC connectivity with the left hippocampus (**Figure. 4b**) and right precuneus (*q* = 0.03, FDR corrected; *Supplementary Table S4*). Next, we observed a negative correlation of child-parent vmPFC-hippocampal functional connectivity with children’s internalizing symptoms, especially for anxious/depressed components (*q* = 0.06, FDR correction).

**Figure 4.**
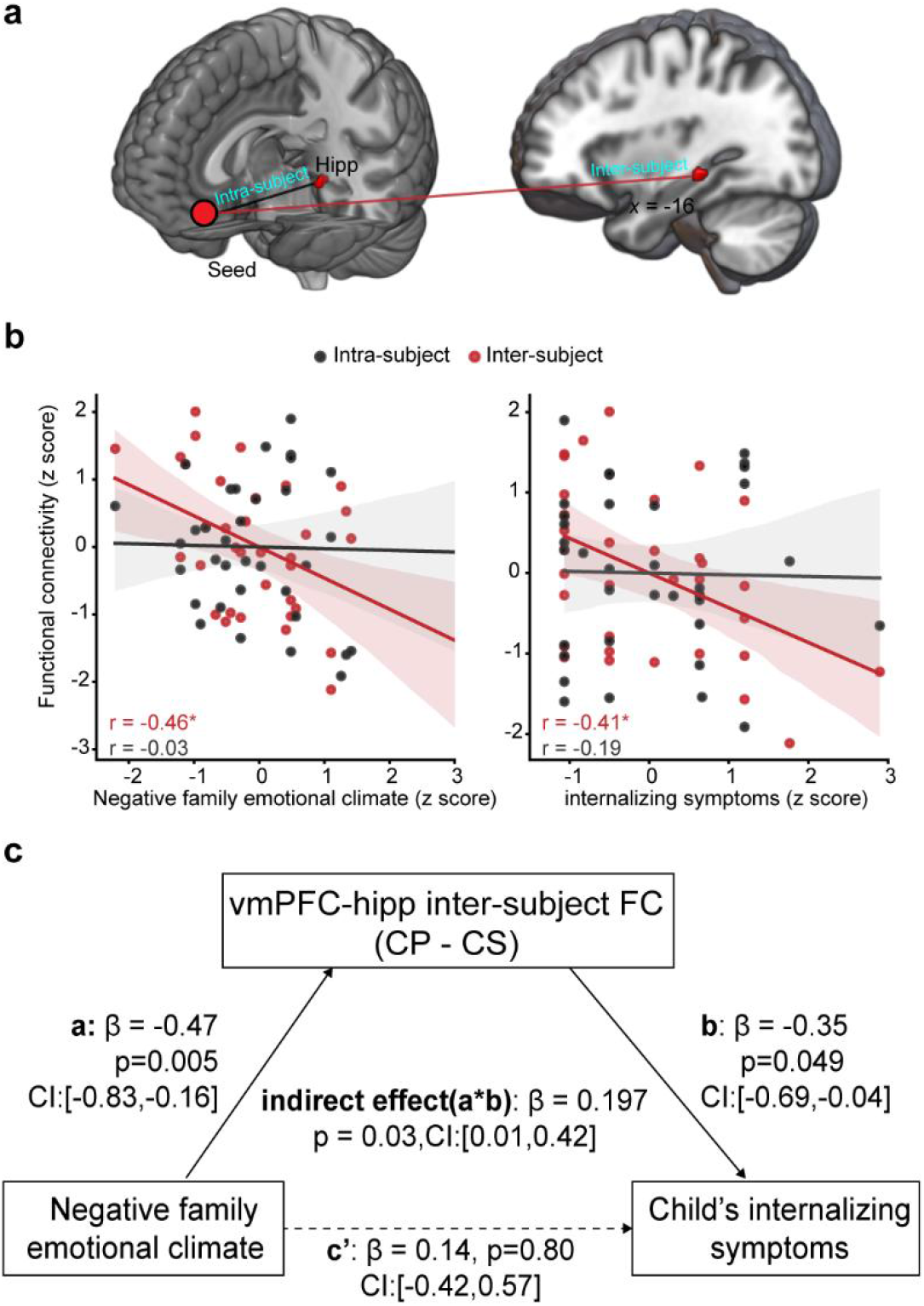
The Relationships between Negative Family Emotional Climate, ISFC and Internalizing Symptoms. *Note*. a. Representative view of a vmPFC seed and its intra-subject and inter-subject connectivity with the hippocampus. b. Scatter plots depict the negative correlations (FDR corrected) of inter-subject vmPFC-hippocampal connectivity (red) with negative family emotional climate and children’s anxious/depressed symptoms. This pattern is not observed using intra-subject vmPFC-hippocampal connectivity (gray). c. A mediation model depicts the indirect pathway of negative family emotional climate on children’s anxious/depressed symptoms via the shared vmPFC-hippocampal inter-subject connectivity. Standardized coefficients are depicted. The solid lines represent statistically significant effect. Notes: \**q* < 0.05. All statistical tests here are two-tailed and pass the FDR correction.

**Figure 5.**
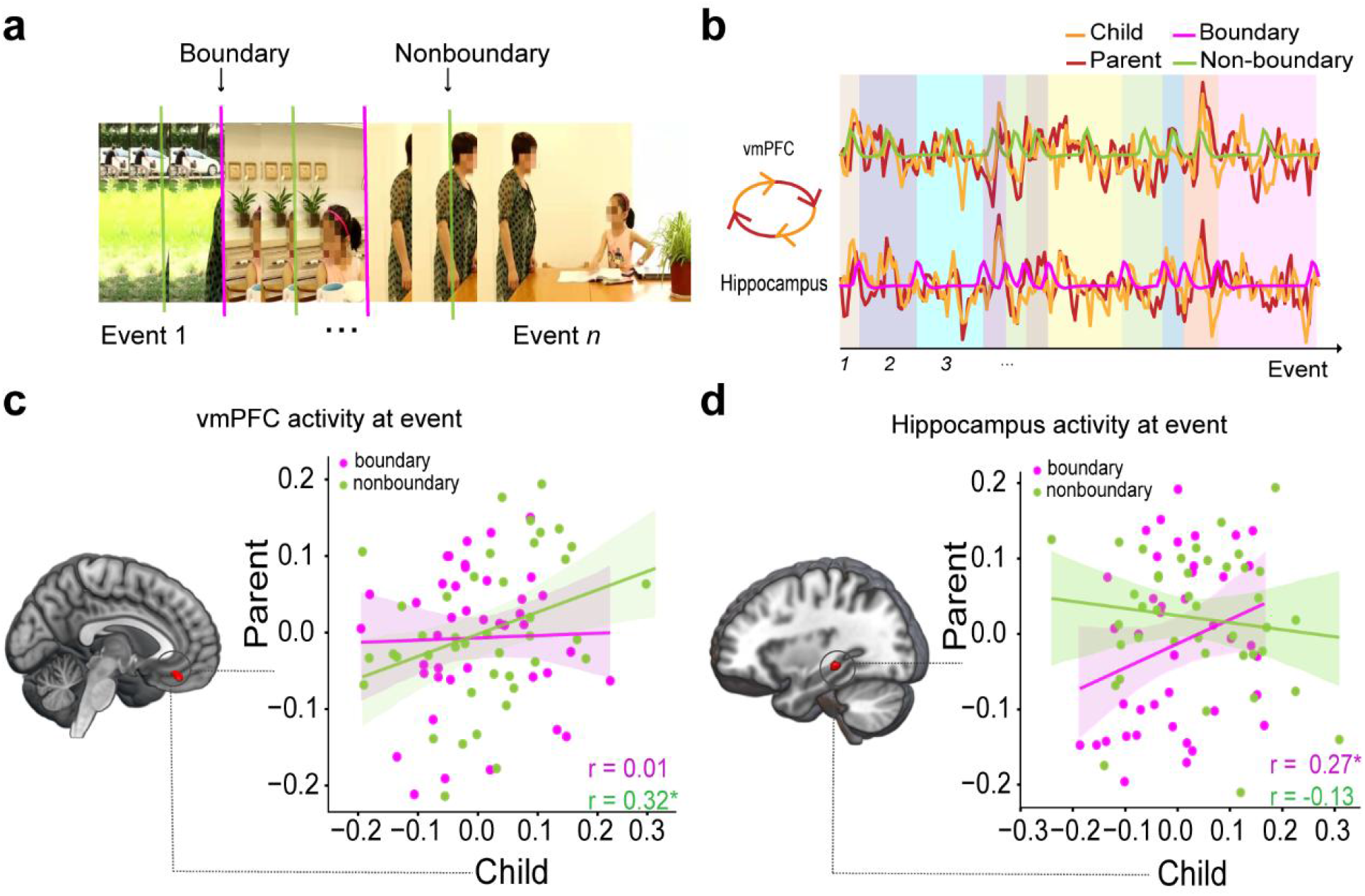
Child-Parent Hippocampal and Vmpfc Activity Concordance during Movie Watching. *Note*. a. An illustration of boundary and non-boundary events for major episodic events during movie watching. b. Child-parent dyads showed higher vmPFC-hippocampus functional coupling during movie watching, and their vmPFC and hippocampal activity concordance were modulated by segmentation of event boundaries. The magenta and green lines represent expected signals of event boundaries and non-boundaries respectively. The yellow and red lines represent neural signals in children and parents separately. c. Child-parent hippocampal activity concordance was significantly higher for boundary than non-boundary event time series (*Z* = 1.84, *p* = 0.03). d. Child-parent vmPFC activity concordance was marginally significant higher for boundary than non-boundary timeseries (*Z* = −1.39, *p* = 0.08).

Since child-parent vmPFC-hippocampal connectivity was associated with both negative family emotional climate and children’s internalizing symptoms, we conducted a mediation analysis to examine whether this inter-subject functional pathway accounts for the association between negative family emotional climate and children’s internalizing problems. This analysis revealed an indirect pathway of reduced vmPFC connectivity with the hippocampus mediating the association between negative family emotional climate and higher children’s internalizing symptoms (**Figure. 4c**, *B* = 0.17, *SE* = 0.10, *p* = 0.028, bootstrapped 95% CI = [0.01,0.42], 56.7% of the total effect size) and children’s anxious/depressed symptoms (*B* = 0.19, *SE* = 0.12, *p* = 0.04, bootstrapped 95% CI = [0.00,0.46], 59.4% of the total effect size). Notably, the mediation effect was significant even when regressing out child-parent’s age and gender (*Supplementary Figure. S4b*). Since children’s emotional problems could influence family emotional climate (Rothenberg et al., 2020), we tested an alternative model with children’s internalizing symptoms as input variable and negative family emotional climate as a outcome predictor. Although this model is also valid (*Supplementary* Figure. S4c&d), model comparison with Bayesian Information Criterion (BIC) favors the initial model with family emotional climate affecting child internalizing symptoms (BIC = 113.89) over the reverse alternative model (BIC = 228.34)(Raftery, 1995).

To verify whether negative family emotional climate is associated with children’s internalizing problem through shared rather than individual vmPFC-hippocampus responses, we performed vmPFC-seeded functional connectivity within children’s brains **(Figure. 4a)**. We did not find any reliable effects pertaining to intra-brain metrics **(Figure. 4b)**. In addition, we conducted time-lagged analysis for vmPFC-based ISFC to determine when child-parent dyads exhibited the highest ISFC. This analysis revealed that child-parent dyads exhibited highest vmPFC-hippocampal functional correlation at lag zero (*Supplementary Figure. S3*). Together, these results indicate that reduced child-parent vmPFC connectivity with the hippocampus accounts for the adverse effects of negative family emotional climate on children’s internalizing symptoms.

### Child-Parent Vmpfc and Hippocampal Activity Concordance in Event Boundaries during Movie Watching

According to recent neurocognitive models of event segregation, the hippocampus and vmPFC are recognized to support segregation of event boundaries and integration of episodic events into structured representations (Baldassano et al., 2017; Ben-Yakov & Henson, 2018; Ezzyat & Davachi, 2021). Our perception and process of such events are actively shaped by existing memories and schematic scripts about experiences in the world(Baldassano et al., 2018). Living in the same family environment, child-parent dyads often have reciprocal interactions during various social and emotional scenarios, and thus they tend to form shared mental schemas on how to understand, cope and respond to key events. Thus, we expected that children would exhibit a similar pattern of hippocampal responses to the moments of shift between meaningful events (boundary timepoints) with their parents.

We therefore implemented dyad-based analysis of brain responses to event segmentation during movie watching to examine whether children’s hippocampal and vmPFC responses to event boundary and non-boundary timepoints are correlated with their parents. As expected, this analysis revealed that children’s hippocampal responses to event boundaries were indeed positively associated with their parent’s responses (*r* = 0.27, *p* = 0.04, 95% CI = [0.04,0.48]). This concordance, however, did not emerge for within-event time points (*r* = −0.13, *p* = 0.21, 95% CI = [-0.32, 0.26]). Further Z-test analysis for two correlations coefficients reveals a significant difference (*Z* = 1.84, *p* = 0.03). Interestingly, parallel analysis revealed an opposite pattern of child-parent concordance for the vmPFC activity. That is, children’s vmPFC responses to non-boundary timepoints were positively correlated with their parents (*r* = 0.33, *p* = 0.02, 95% CI = [0.03, 0.54]), but not for event boundaries (*r* = 0.03, *p* = 0.42, 95% CI = [-0.22, 0.30]). Further tests revealed a marginally significant difference between the two correlations (*Z* = −1.39, *p* = 0.08). Taken together, these results indicate that the vmPFC and hippocampus exhibit interactive activity concordance in child-parent dyads in response to non-boundary and boundary events during movie watching.

## Discussion

By leveraging dyad-based analysis of fMRI data during naturalistic movie watching, we investigated the neural substrates of how negative family emotional climate was associated with children’s psychopathological symptoms through child-parent brain-to-brain activity and connectivity concordance. Compared to child-stranger dyads, child-parent dyads exhibited higher inter-subject correlation in the vmPFC and dmPFC, and higher inter-subject connectivity of the vmPFC with widespread regions critical for socioemotional cognition. Critically, reduced child-parent vmPFC-hippocampal connectivity accounted for the association between negative family emotional climate and children’s internalizing symptoms, with distinct contributions of the vmPFC and hippocampus to non-boundary and boundary events, respectively, during movie watching. Our findings provide a neurobehavioral model of how negative family emotional climate is associated with children’s internalizing symptoms through reduced child-parent brain-to-brain concordance in the vmPFC-hippocampal circuitry.

Behaviorally, children in negative family emotional climate experienced more severe psychopathological symptoms as indicated by heightened internalizing and externalizing problems. This is in line with previous findings showing positive associations between family risk factors (e.g., maternal maltreatment, family conflicts) and children’s internalizing and externalizing symptoms (Gong et al., 2021; Schleider & Weisz, 2017),. According to social learning and bio-behavioral synchrony models (Feldman, 2020a; Justyna, 2017), child-parent shared experiences are indispensable for children to learn and develop emotional skills through socialization from their parents in daily life. Through child-parent reciprocal interactions such as affective synchrony and empathic dialogues, for instance, children regulate themselves to attune to each other’s minds, which helps them develop socioemotional skills such as emotion regulation and theory of mindand then reduce the risk of suffering psychopathological symptoms (Feldman, 2020; Thomassin & Suveg, 2014]).

Socioemotional interactions in a family have also been demonstrated to help child-parent dyads build a shared or synchronous pattern of brain responses (Piazza et al., 2020; Wass et al., 2020). Such shared neural responses can serve as a scaffold for children’s socialization helping children learn necessary skills and form memories through reciprocal interactions with their parents in daily life (Reindl et al., 2018b). Thus, negative family environment (e.g., negative family emotional climate) may impede child-parent brains from forming effective socioemotional skills, accounting for the emergence of children’s psychopathological symptoms. As discussed below, this account is supported by three aspects of our observed concordance across child-parent brains.

Our movie-watching fMRI results show that children-parent dyads exhibited higher inter-subject correlation in the vmPFC and dmPFC than control child-stranger dyads. This is reminiscent of previous findings showing that the mPFC plays a critical role in characterizing shared neurocognitive processes between children and their parents (Hoyniak et al., 2021a; Itahashi et al., 2020; Piazza et al., 2020). The mPFC is thought to act as a simulator for social event schemas that allows us to integrate and summarize social, self and emotional information as events unfold over time (Krueger et al., 2009a). When processing socioemotional events, the dmPFC is important for inferring other’s goal-oriented actions, whereas the vmPFC is crucial for appraisal, evaluation and regulation of values involved in self and affective processes (Bzdok et al., 2013). Such processes could serve as a neurocognitive basis to understand the intentions and mental states of others (Fiske & Taylor, 2013). Thus, higher inter-subject correlation in the dmPFC and vmPFC across child-parent dyads likely reflect similarities in how child-parent dyads perceive and react emotionally to the socioemotional world.

Our results also show that child-parent dyads exhibited higher inter-subject correlation of vmPFC- and dmPFC-based functional connectivity with widespread regions of social and emotional brain networks in comparison with child-stranger dyads. Specifically, child-parent dyads shared vmPFC coupling with distributed regions crucial for episodic memory, emotion and social processing (Lieberman et al., 2019; Phillips et al., 2019), while shared dmPFC coupling had relatively uniform connectivity with regions such as TPJ during movie watching. These inferences were drawn from a widely used reverse inference database (Yarkoni et al., 2011). The vmPFC and its coordination with the hippocampus, precuneus, amygdala and among others are recognized to support the appraisal of perceived socioemotional events (Hiser & Koenigs, 2018) and the reinstatement of existing knowledge and strategies formed over the course of child-parent interactions (Feldman, 2015, 2017). These processes help children learn how to cope with negative emotions(Nawa & Ando, 2019; Roy et al., 2012b). Our data suggest that vmPFC circuitry is critical for integration of disparate events shared by child-parent dyads when viewing emotionally negative movies, likely by promoting transmission of affectivity and sociality across child-parent dyads.

Furthermore, our fMRI results additionally showed that reduced brain-to-brain concordance as indicated in the vmPFC-hippocampal pathway in child-parent dyads mediated the association between negative family emotional climate and more severe child internalizing symptoms, highlighting a possible mechanism for how negative family emotional climate associates with children’s psychopathological symptoms. This finding provided one of the first empirical evidence of the Extended Parent–Child Emotion Regulation Dynamics Model (Ratliff et al., 2022), supporting that cross-brain connectivity between child and parent serves as an important mechanism linking family environment with child emotional development, and the vmPFC-hippocampal circuitry is important for constructing the meaning of emotional events (Nawa & Ando, 2019; Roy et al., 2012b). Child-parent concordance of vmPFC-hippocampal coupling during movie-watching likely reflects their co-construction of socioemotional events according to shared and/or embodied relationships. Constructing the meaning of emotional events can be a useful way to regulate children’s internalizing states to achieve psychological balance (Carpendale & Lewis, 2004). It is possible that children with higher parental concordance of vmPFC-hippocampal connectivity may develop better socioemotional skills and thus exhibit lower levels of internalizing symptoms. Our observed mediation effect suggests that child-parent vmPFC-hippocampal concordance could serve as a potential biomarker for children in families with emotional disorders. Future work may use neurofeedback techniques to explore the impact of upregulating vmPFC-hippocampus coordination with parents on children’s emotional health.

VmPFC-hippocampal circuitry is also crucial in updating and integrating new events into existing memory schemas(Gilboa & Marlatte, 2017; Zeithamova et al., 2012). Increased vmPFC-hippocampal concordance may reflect child-parent dyads with lower negative family emotional climate by updating their internal schemas when socioemotional events are incongruent with existing schemas. Although we cannot rule out this possibility, analysis of event boundary-evoked response revealed that children’s hippocampal responses to event boundaries were positively related to their parent’s responses. Given that segmenting continuous events into meaning units is driven by our experience and mental schemas (Baldassano et al., 2018), child-parent hippocampal activity concordance on boundary-evoked response suggests that child-parent dyads utilize their shared episodic memories and schemas to understand and interpret socioemotional events during movie watching. Together with stronger inter-subject vmPFC-hippocampal connectivity observed in child-parent dyads, our results are among the first to suggest that the vmPFC may signal child-parent hippocampal activity concordance in order to orchestrate long-term memory, emotional and social systems in support of their understanding of events during movie watching.

Several limitations should be considered in our study. First, we assessed child-parent neural concordance at activity and connectivity levels when viewing emotionally negative movies. Whether our findings can be generalized into positively valenced situations still remains open for future studies. Second, although we leveraged a naturalistic movie-watching fMRI paradigm, dedicated task designs are needed to complement the interpretation of child-parent shared neural responses in vmPFC and related circuits. Finally, it is worth noting that we also observed the indirect effect of child-parent shared brain responses in the association of negative family emotional climate with children’s internalizing problems. It is thus possible that such relationships are bidirectional (Gong et al., 2021; Nelemans et al., 2020). Longitudinal designs are required to disentangle the directionality effects.

## Conclusion

The present study demonstrates brain-to-brain concordance across child-parent dyads during movie watching that was localized to the mPFC and its connectivity with regions in socioemotional networks. Inter-brain concordance in ventral mPFC-hippocampal circuitry, rather than intra-brain metrics, emerged as a key locus that mediates the adverse effect of negative family environment on children’s internalizing symptoms. Our study provides a neurobiological account for how negative family environment influences children’s internalizing symptoms through shared socioemotional representations across brains in child-parent dyads. This work can inform the development of dyad-based prevention and interventions designed to improve children’s internalizing problem.

## Author contributions

S.Q. and Z.R.H. conceptualized this project. H.S. collected data. J.W. contributed to cleaning data. H.S. and G.C. performed data analyses. H.S., C.B.Y., Z.R.H., J.X. and S.Q. wrote the manuscript.

## Competing Interests

The authors declare that they have no competing interests.

## Data and Materials Availability

All data needed to evaluate the conclusions er are present in the paper and/or the Supplementary Materials. Raw data may be requested from the corresponding authors.

## Notes

### Competing Interest Statement

The authors have declared no competing interest.

### Summary of Updates

Supplemental files updated.

