## Supplementary Information for "Atypical Child-Parent Neural Synchrony Links to Children’s Psychopathological Symptoms"

Supplementary Figures Legends S1-S4

**Supplementary Figure. S1.**

*The Partial Correlation between Family Emotional Climate and Children's Behavioral Problems.*

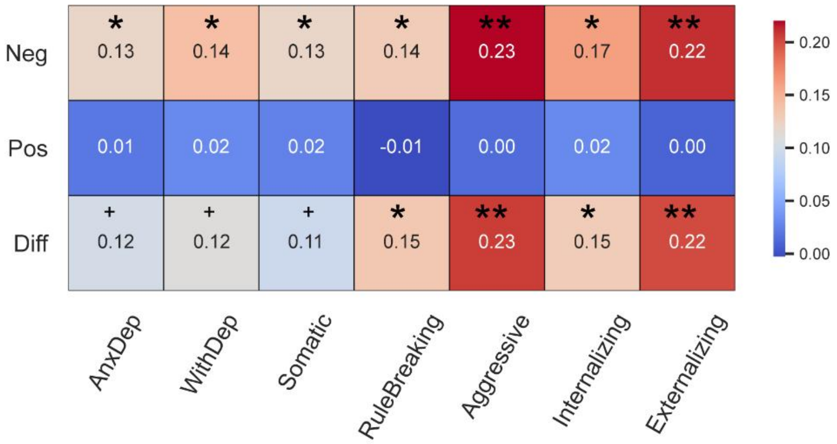


*Note.* Partial correlations of negative (row 1) and positive (row 2) family emotional climate with children's anxious/depressed, withdrawn/depressed, somatic, rule breaking, aggressive, internalizing and externalizing problem scores after controlling for children's age, gender, parent's gender and family socioeconomical status. The absolute differences between row1 and row2 are shown in row 3. (FDR correction, +$q$ = 0.06, * $q < 0.05$, ** $q < 0.01$)

**Supplementary Fig. S2.**

*Results from Dmpfc-Seed Functional Connectivity (ISFC) Analysis.*

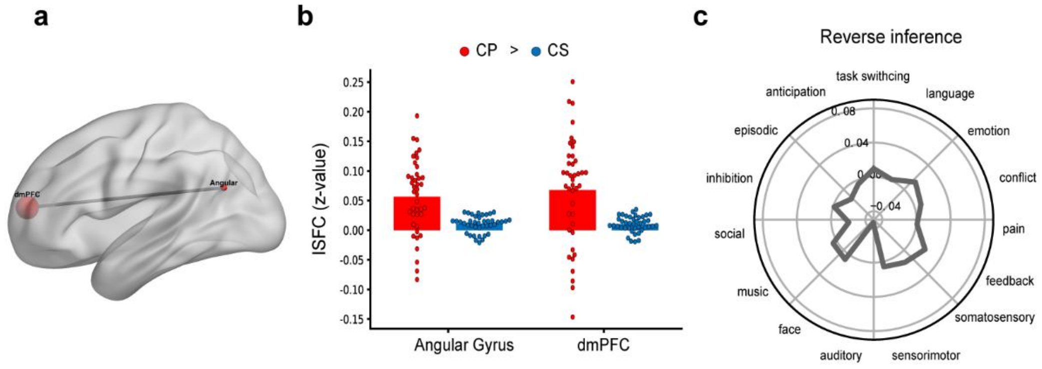


*Note.* a. Child-parent dyads were more synchronous than the child-stranger dyads between dmPFC, angular gyrus and the voxels surrounding the dmPFC (FDR correction, $q$ value < 0.05). b. The ISFC thresholded map based on the contrast between child-parent and child-stranger dyads revealed a uniform association with the psychological domains.

**Supplementary Fig. S3.**

*The Time-Lagged Functional Correlation.*

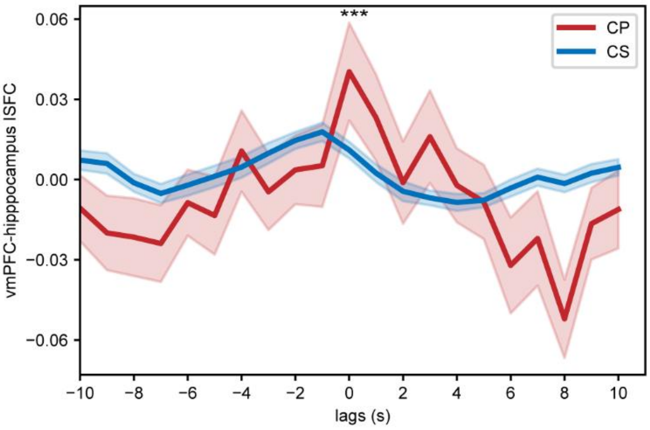


*Note.* The time-lagged functional correlation among pairs showed child-parent dyads had the highest shared vmPFC-hippocampus functional connectivity at the time 0.

**Supplementary Figure. S4.**

*The Relationships between Internalizing Symptoms, ISFC and Negative Family Emotional Climate.*

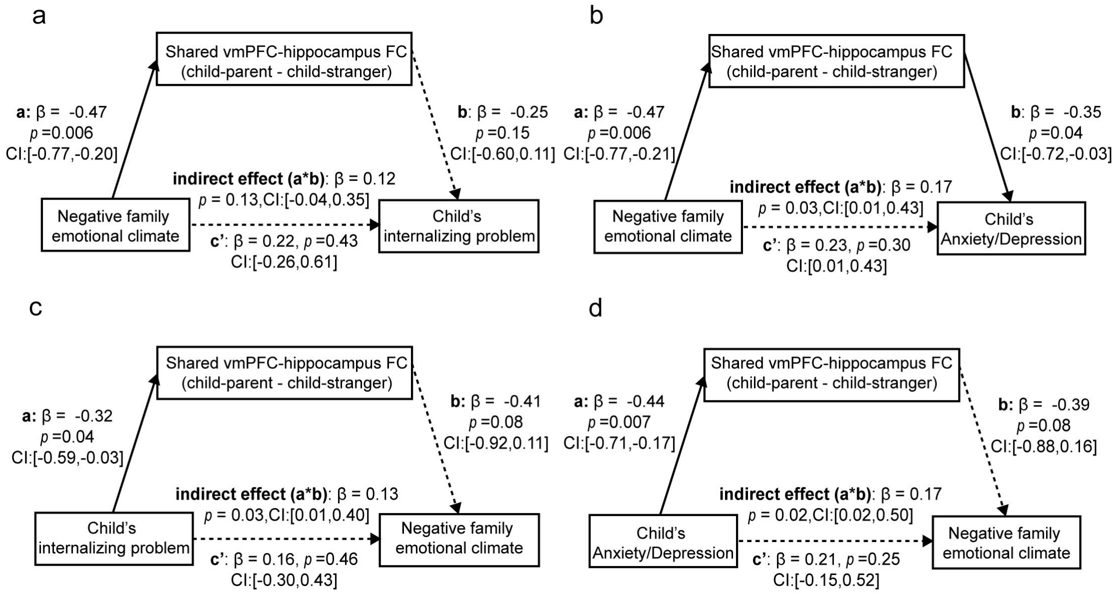


*Note.* Mediation models depict the relationships between negative family emotional climate, shared vmPFC-hippocampus functional connectivity, and children's internalizing as well as anxious depressed problem scores after control child's gender, age, parent gender and site (N=35). **a.** Shared vmPFC-hippocampus functional connectivity did not significantly mediate the association between negative family emotional climate and children's internalizing problems (B = 0.12, SE = 0.10, p = 0.13, bootstrapped 95% CI = [-0.04,0.35], 35.3% of the total effect size). **b.** Shared vmPFC-hippocampal inter-subject connectivity significantly mediated the association between family emotional climate and children's anxious/depressed problems (B = 0.17, SE = 0.10, $p$ =0.023, bootstrapped 95% CI = [0.02,0.49], 42.5% of the total effect size). **c.** Shared vmPFC-hippocampal inter-subject connectivity significantly mediated the association between children's internalizing problems and negative family emotional climate (B = 0.13, SE = 0.09, $p$ =0.03, bootstrapped 95% CI = [0.01,0.40], 44.8% of the total effect size). **d.** Shared vmPFC-hippocampal inter-subject connectivity significantly mediated the association between children's anxious/depressed problems and negative family emotional climate (B = 0.17, SE = 0.11, $p$ = 0.02, bootstrapped 95% CI = [0.02, 0.50], 44.7% of the total effect size). Standardized coefficients are depicted. The solid lines represent significant effect. Notes: *$p < 0.05$; **$p < 0.01$. All mediation effect tests here are two-sided, and pass the FDR correction.

Supplementary Tables S1-S4

**Supplementary Table S1.**

*The Critical Moments of the Parent-Child Conflict Video Used for the Fmri Paradigm.*

| | Time | Major movie events |
|---|---|---|
| 1 | 00:00-00:10(10 s) | A girl and her mother sit on chairs after crossing the forest path. |
| 2 | 00:10-00:50(39s) | The mother scolds the girl. The girl is nervous and wants to cry. |
| 3 | 00:51-01:33(42s) | The child stands up excitedly, shouts, and argues with her mother. |
| 4 | 01:34-01:48(14s) | The mother shows impatience with the child and turns her head. The girl sits down with her back to her mother. |
| 5 | 01:49-02:00(11s) | At lunch, the girl, fed up with her mother's scolding, puts her hands over her ears. |
| 6 | 02:01-02:16(15s) | Both the girl and the mother throw down their chopsticks. |
| 7 | 02:17-03:07(50s) | The mother keeps slapping the table and criticizing the child. |
| 8 | 03:08-03:32(25s) | The child turns her head, and her mother slaps the table to make her look back. They argue furiously. |
| 9 | 03:33-03:45(12s) | Seeing the child playing on the computer, the |

| | | |
|---|---|---|
| | | mother wrings her hands and sighs. |
| 10 | 03:46-04:07(22s) | The mother turns off the child's computer. The child stands up and the two argue. |
| 11 | 04:08-05:10(22s) | The mom and girl continue arguing about homework |

**Supplementary Table S2.**

*Characteristics of Participants with and without Brain Imaging Data*

| | **without brain imaging data(n=360)** | | **with brain imaging data(n=35)** | | | |
|---|---|---|---|---|---|---|
| **Variable** | ***M*** | ***SD*** | ***M*** | ***SD*** | ***t(393)*** | ***p*** |
| Negative family emotional climate | 3.40 | 1.21 | 3.81 | 1.10 | 1.89 | 0.06 |
| Positive family emotional climate | 6.32 | 1.37 | 6.5 | 1.16 | 0.69 | 0.48 |
| Internalizing problems | 3.57 | 3.84 | 4.70 | 3.80 | 1.64 | 0.10 |
| Externalizing problems | 4.40 | 4.46 | 4.87 | 3.31 | -0.68 | 0.50 |
| Withdrawn/Depressed | 0.99 | 1.29 | 1.38 | 1.85 | 1.66 | 0.10 |
| Anxious/Depressed | 1.44 | 1.83 | 1.88 | 1.80 | 1.38 | 0.17 |
| Somatic | 1.12 | 1.54 | 1.38 | 1.85 | 0.91 | 0.36 |
| Rule Breaking | 1.26 | 1.45 | 1.44 | 1.43 | 0.72 | 0.47 |
| Aggressive | 3.15 | 3.34 | 3.43 | 2.55 | 0.49 | 0.67 |
| Child age | 9.28 | 1.64 | 10.13 | 1.44 | 2.93 | 0.004 |
| Family socioeconomical status | -0.007 | 0.74 | 0.07 | 0.59 | 0.57 | 0.57 |
| | *n* | *%* | *n* | *%* | ***$\chi^2$(1)*** | *p* |
| Child gender,male | 180 | 0.50 | 19 | 0.54 | 0.22 | 0.64 |
| Parent gender,male | 118 | 0.33 | 8 | 0.22 | 1.47 | 0.23 |

**Supplementary Table S3.**

*Results from Inter-Subject Correlation (ISC) Analysis*.

| Threshold | Contrast | Region | L/R | *T* | MNI (xyz) | K |
|---|---|---|---|---|---|---|
| $p<0.001^{a}$ | child-parent> child-stranger | vmPFC | R | 5.07 | -2,38, -18 | 116 |
| | | dmPFC | L | 6.03 | 0, 56, 12 | 122 |

*Note.* The brain regions showing inter-subject correlation between child-parent dyads and child-stranger dyads. (the initial *p*-value < 0.001 (two-tailed), cluster correction α < 0.05, cluster size=82 voxels)

**Supplementary Table S4.**

*Vmpfc- and Dmpfc-Based Inter-Subject Functional Correlation.*

| Contrast | Region | L/R | *T* | MNI (x y z) | K | Corr[a] |
|---|---|---|---|---|---|---|
| vmPFC: child-parent> child-stranger | Amygdala | R | *5.45* | 18 2 -16 | 76 | -0.20 |
| | Medial Frontal Gyrus | L | *5.17* | -4 40 -20 | 86 | -0.04 |
| | Caudate | L | *6.16* | -4 10 -12 | 54 | -0.20 |
| | Medial Frontal Gyrus | L | *5.05* | -6 46 -10 | 41 | -0.04 |
| | Fusiform Gyrus | L | *5.38* | -32 -48 -8 | 36 | -0.21 |
| | Hippocampus | L | *5.28* | -16 -30 -8 | 31 | -0.46* |
| | Precuneus | L | 6.54 | -12 -56 28 | 276 | -0.35 |
| | Precuneus | R | 5.45 | 20 -52 16 | 35 | -0.49* |
| | Precuneus | R | 5.91 | 22 -60 26 | 52 | -0.31 |
| | Middle occipital gyrus | L | 5.96 | -34 -80 34 | 92 | -0.23 |
| | Superior Frontal Gyrus | L | 5.56 | -26 58 34 | 40 | -0.09 |
| | Middle cingulum cortex | R | 5.13 | 10 -32 32 | 43 | -0.15 |
| | Middle cingulum cortex | L | 5.25 | -8 -26 46 | 72 | -0.34 |
| dmPFC: child-parent> child-stranger | Medial Frontal Gyrus | R | 5.75 | 0 54 14 | 37 | 0.01 |
| | Angular | L | 5.8 | -46 -64 24 | 36 | -0.16 |

*Note.* The brain regions showing increased vmPFC- and dmPFC-based inter-subject functional correlation between child-parent and child-stranger dyads (FDR correction, $q<0.05$). a. The correlation between each cluster and negative family emotional climate (FDR correction, *$q$ value<0.05).
